# Transcriptomics-Conditioned Virtual Tissue Synthesis via Diffusion Transformers

**DOI:** 10.64898/2026.05.26.727902

**Authors:** Pantelis R. Vlachas, Kalin Nonchev, Viktor H. Koelzer, Gunnar Rätsch

**Author notes:** Correspondence to: Pantelis R. Vlachas < >, Kalin Nonchev < >, Viktor H. Koelzer < >, Gunnar Rätsch < >. Equal contribution (Pantelis R. Vlachas, Kalin Nonchev). Equal supervision (Viktor H. Koelzer, Gunnar Rätsch).

## Abstract

Spatial transcriptomics couples hematoxylin and eosin (H&E) tissue morphology with spatially resolved gene expression (GE). However, generative models that exploit this coupling to synthesize tissue images from transcriptomic profiles remain scarce. We present STMDiT (Spatial Transcriptomics and Morphology Diffusion Transformer), a diffusion transformer that synthesizes H&E histopathology patches conditioned jointly on morphological embeddings and transcriptomic profiles. Building on PixCell (Yellapragada et al., 2025), we integrate gene expression from a frozen CancerFoundation encoder (Theus et al., 2024) through adaptive layer normalization and per-block cross-attention, and we train under dual classifier-free guidance with independent modality dropout. On the 10x TuPro Visium melanoma cohort, GE conditioning improves both image quality over the no-GE PixCell-B baseline (best FID = 252.9 vs 330.7) and transcriptomic fidelity (best AUC = 0.267 vs 0.229, reaching 82% of the real-tile ceiling). Training with DeepSpot’s predicted-transcriptomics pseudo-labels (PTPL) uniquely transfers zero-shot to TCGA SKCM, an out-of-distribution (OOD) H&E-only melanoma cohort: PTPL-XAttn-PMA-B reaches FID = 690.0, a 57-point improvement over the no-GE baseline (747.1), with a within-model GE-ablation effect of Δ_OOD_ = +309.5, enabling virtual tissue synthesis beyond native spatial-transcriptomics coverage. Our results indicate that gene-expression conditioning produces morphologically distinct tissue images and supports virtual tissue simulation for hypothesis testing in computational pathology.

## 1. Introduction

Spatial transcriptomics (ST) platforms such as 10x Visium, MERFISH, and Slide-seq measure gene expression and tissue morphology in the same section at spot or singlecell resolution (Ståhl et al., 2016; Williams et al., 2022), pairing an H&E image with the gene expression of the cells within each captured region. This pairing supports tumor microenvironment analyses (Yoosuf et al., 2020) but current ST platforms remain costly, cover limited tissue areas, and are not routinely used in clinical diagnostics (Choe et al., 2023).

Prior work spans two complementary directions. Diffusion models for pathology have progressed from text-conditioned latent diffusion on pathology reports (Yellapragada et al., 2024) to foundation models such as PixCell (Yellapragada et al., 2025) (30M H&E patches with UNI2-h morphology conditioning (Mahmood Lab, 2024)) and CytoSyn (Duboudin et al., 2026), all conditioning on morphology or text alone. In the inverse direction, H&E → GE predictors regress spot expression directly from tissue patches: HE2RNA (Schmauch et al., 2020), HisToGene (Pang et al., 2021), BLEEP (Xie et al., 2023), DeepSpot (Nonchev et al., 2025) (state-of-the-art on the 10x TuPro melanoma cohort used in this work), and the recent STORM foundation model scaling to 1.2M spots (Xiang et al., 2026).

Generating tissue morphology jointly from a reference image and a transcriptomic profile would enable virtual tissue experiments: perturbing gene-expression inputs and observing how tissue appearance changes, or augmenting datasets with controlled transcriptomic variation while preserving spatial context. Pixel-aligned conditioning such as ControlNet (Zhang et al., 2023) does not apply directly, since a Visium spot aggregates 1 to 10 cells into a single expression vector without spatial substructure aligned with the image. Several concurrent works synthesize images directly from gene expression. Wu et al. (2024) edit gene-expression values to generate gigapixel mouse-pup images via a GAN. In parallel, Wu et al. (2025) synthesize 3D mouse-brain volumes from spatial mRNA via diffusion. On H&E, GeneFlow (Wang et al., 2025) produces tiles from Xenium single-cell expression via a rectified-flow U-Net, and Lohmann et al. (2025) contrastively align gene expression with UNI embeddings to condition a GAN or fine-tuned Stable Diffusion. All four use gene expression as the sole conditioning signal at inference, with no mechanism to additionally condition on a reference morphology.

We introduce STMDiT^1^, a dual-conditioned diffusion transformer that extends PixCell (Yellapragada et al., 2025) with a gene-expression stream from a frozen CancerFoundation encoder (Theus et al., 2024). We compare two fusion architectures, adaptive layer normalization (AdaLN) and per-block cross-attention, and evaluate whether the generated images preserve and reflect the transcriptomic conditioning signal through downstream regression and a tissue-level gene expression (GE)-perturbation protocol. The PTPL variant replaces raw Visium counts with DeepSpot’s (Nonchev et al., 2025) denoised gene-expression predictions, enabling zero-shot extension to H&E-only cohorts such as TCGA SKCM.

## 2. Method

### 2.1 Architecture and Gene Expression Conditioning

Our backbone follows PixCell (Yellapragada et al., 2025). An SD3 variational autoencoder (VAE) (Esser et al., 2024) encodes each 256 × 256 H&E patch into a latent *z* ∈ ℝ^32*×*32*×*16^, which a PixArt-Σ diffusion transformer (Peebles & Xie, 2023; Chen et al., 2024b;a; Rombach et al., 2022) denoises under UNI2-h cross-attention (Chen et al., 2024c; Mahmood Lab, 2024). We train either with the standard denoising objective over *T* = 1000 timesteps or with a flow-matching objective (Lipman et al., 2023; Esser et al., 2024) in the flow variants.

We additionally condition the diffusion model on a geneexpression (GE) profile 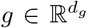 extracted from a frozen CancerFoundation encoder (Theus et al., 2024), a transformer scRNA-seq foundation model pretrained on malignant cells whose latent representation captures cross-gene regulatory structure. We investigate two fusion architectures, illustrated at the block level in Figure 2 (Section B).

#### AdaLN fusion

The GE embedding is projected to the timestep-embedding dimension and combined with the timestep embedding through a learnable gate. The fused embedding modulates each transformer block via adaptive layer normalization, following the standard PixArt conditioning pathway (Chen et al., 2024b). No new attention layers are introduced, making this variant parameter-efficient.

#### Cross-attention fusion

The GE embedding is transformed into a sequence of *K* tokens that are attended to via per-block cross-attention alongside the UNI embedding. We evaluate four strategies for producing the token sequence: (i) *Direct*, the raw CancerFoundation output (roughly 1200 tokens); (ii) *PMA*, pooling by multi-head attention with learnable query vectors to *K*=32 tokens (Lee et al., 2019); (iii) *Perceiver*, iterative cross-attention with learnable latents (*K*=32) (Jaegle et al., 2021); (iv) *GSA*, self-attention over gene tokens (*K*=32).

### 2.2 Dual Classifier-Free Guidance

During training, we independently drop each conditioning modality with probability *p*, following the standard classifier-free-guidance recipe (Ho & Salimans, 2021). This yields four regimes (full conditioning, UNI-only, GE-only, unconditional) with training weights (1 − *p*)^2^ : *p*(1 − *p*) : *p*(1 − *p*) : *p*^2^. At inference we apply separate guidance scales *s*_uni_ and *s*_ge_:

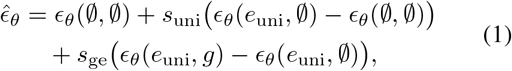

where ∅ denotes a null token, and *ϵ*_*θ*_(∅, *g*) is trained but unused at inference. The dropout probability *p* acts as a modality-balancing knob: low *p* leaves UNI dominant over the GE stream, while larger *p* rebalances by increasing the fraction of GE-only and unconditional samples. The exact *p* values trained per family are listed in Section 3.

## 3. Experimental Setup

### Dataset

We train and evaluate on the Tumor Profiler (TuPro) dataset (Nonchev et al., 2024): 18 metastatic melanoma slices from 7 donors on 10x Visium at ∼55 *µ*m spots covering 1 to 10 cells each, with pathologist-reviewed tissue labels used only for the tissue-level perturbation analysis (Section K) and the per-tissue quality breakdown (Section J). Main-text FID and AUC are spot-level and label-agnostic. We split at the donor level, holding out one patient for evaluation. For each spot we pre-extract an SD3-VAE latent (Esser et al., 2024), a UNI2-h morphological embedding (Chen et al., 2024c; Mahmood Lab, 2024), and a Cancer-Foundation GE embedding (Theus et al., 2024). The PTPL variant replaces the GE *conditioning input* with a DeepSpot (Nonchev et al., 2025) prediction, denoising raw counts via spatial aggregation across neighboring spots; the split, UNI embeddings, and VAE latents are unchanged. Spot counts, filtering, and hyperparameters are in Sections A and D.

### Models

All trained models use the PixArt-B architecture (hidden 768, depth 12). We train two no-GE baselines (PixCell-B and PixCell-Flow-B), AdaLN DDPM/Flow and cross-attention (Direct, PMA, Perceiver, GSA) GE variants across a modality-dropout sweep *p* ∈ {0.1, 0.2, 0.3, 0.5, 0.6}, and matched PTPL counterparts for three architectures at *p* ∈ {0.5, 0.6, 0.7}. PixCell-Pretrained-XL (Yellapragada et al., 2025) serves as an external pan-cancer reference; full per-family dropout enumeration in Section C.

### Evaluation

We report FID on H0-mini features (Saillard et al., 2024) across the four conditioning regimes (UNI+GE, UNI, GE, unconditional), and transcriptomic fidelity via two independent H&E → GE regressors trained on real (image, GE) pairs: an *Oracle* (UNI2-h with principal component analysis (PCA) and Ridge regression on top-*k* highly variable genes (HVGs), inspired by HEST (Jaume et al., 2024; Chen et al., 2024c; Mahmood Lab, 2024)) and a *Mid-night+MLP* (multi-layer perceptron) head on all 19,338 genes (Karasikov et al., 2025). Each regressor is summarized by the area under its per-gene-count Pearson *r* curve (*k* ∈ {50, 100, 200, 500, 1000}), with a real-tile upper bound and a zero-correlation null as anchors. We additionally report a tissue-level GE-perturbation composite: we interpolate the input GE embedding between two tissue-class centroids in CancerFoundation space and measure the H0-mini distance moved toward the target class. Since observational ST provides no paired counterfactual, this composite is a distance proxy rather than a biological-validity claim. Confidence intervals use a common spot-level bootstrap (*n*=11,776, 1000 resamples). Full protocols in Section D

## 4. Results

### 4.1. Image Quality

Table 1 reports test-split FID across the four conditioning regimes for best-of-family models on 10x TuPro. The in-domain no-GE baselines sit at FID=330.7 (PixCell-B) and FID=342.5 (PixCell-Flow-B), and the pretrained pan-cancer XL baseline reaches FID=298.9. Every GE-conditioned family beats both in-domain baselines at its best configuration: AdaLN DDPM reaches FID=274.1 at *p*=0.6, AdaLN Flow reaches FID=293.5 at *p*=0.5, and cross-attention reaches the overall best FID=**252.9** (Perceiver, *p*=0.5).

**Table 1.** Test FID (H0-mini, lower is better) across conditioning regimes for best-of-family models on 10x TuPro. Δ = FID(UNI+GE) *−* FID(UNI); negative values mean GE improves image quality. UNI+GE values are bootstrap means at *n*=11,776; 95% CIs are in Section F.

| Model | $p$ | UNI+GE | UNI | GE | $\Delta$ |
| --- | --- | --- | --- | --- | --- |
| PixCell-Pretrained-XL | – | – | 298.9 | – | – |
| PixCell-B | – | – | 330.7 | – | – |
| PixCell-Flow-B | – | – | 342.5 | – | – |
| AdaLN DDPM (best) | 0.6 | **274.1** | 268.8 | 446.0 | +5.4 |
| AdaLN Flow (best) | 0.5 | 293.5 | 293.4 | 508.6 | +0.1 |
| XAttn-Direct | 0.1 | 285.3 | 497.9 | 602.5 | **−212.6** |
| XAttn-Perceiver | 0.5 | **252.9** | 255.7 | 431.4 | −2.8 |
| XAttn-PMA | 0.5 | 256.7 | 436.8 | 423.0 | −180.1 |

The conditioning ablation exposes how much each family actually uses GE at inference. AdaLN DDPM is almost GE-insensitive across the full dropout sweep (Table 4). Removing GE changes FID by at most 5.7 points at any *p* ∈ {0.1, 0.2, 0.3, 0.5, 0.6}, and reverting to GE-only collapses the generator to FID ≥ 446. GE removal collapses XAttn-Direct (Δ= − 212.6) and XAttn-PMA (Δ= −180.1) but only nudges XAttn-Perceiver (Δ= −2.8): cross-attention splits by token source.

### 4.2 Gene Expression Preservation

Table 2 reports integrated AUC under the UNI2-h+PCA+Ridge Oracle and the Midnight+MLP regressor. Both regressors agree on the direction: every GE-conditioned model beats the no-GE PixCell-B baseline on Oracle AUC (0.229) and on Midnight+MLP AUC (0.118). They disagree on the headline model. Oracle ranks XAttn-Direct highest at 0.267 against a real-tile upper bound of 0.325, while Midnight+MLP places AdaLN Flow at the top (0.131) essentially at the real-tile ceiling (0.130), with the other GE-conditioned rows within 0.005 of it. Across the GE-conditioned rows of Table 2 the two regressors disagree on fine-grained ordering while agreeing on direction, so the overall finding of GE preservation is regressor-agnostic.

**Table 2.** Integrated AUC under two H&E *→* GE regressors (higher is better): the UNI2-h+PCA+Ridge Oracle on top-*k* HVGs and the Midnight+MLP regressor on all 19,338 genes. Both regressors are trained on real (image, GE) pairs from the same 10x TuPro train split and evaluated against real Visium ground truth on the test split. Values are shown as AUC *±* CI (% of real-tile ceiling in parentheses), where CI is the 95% spot-bootstrap half-width (*n*=11,776, 1000 resamples). Percentages use the mean ceiling as denominator.

| Model | Oracle AUC (% ceiling) | Midnight+MLP AUC (% ceiling) |
| --- | --- | --- |
| Upper bound (real tiles) | 0.325 ± 0.002 (100%) | 0.130 ± 0.002 (100%) |
| Null baseline | 0.000 (0%) | 0.000 (0%) |
| PixCell-Pretrained-XL | 0.223 ± 0.002 (69%) | 0.092 ± 0.002 (70%) |
| PixCell-B | 0.229 ± 0.002 (70%) | 0.118 ± 0.002 (91%) |
| PixCell-Flow-B | 0.226 ± 0.002 (70%) | 0.123 ± 0.002 (94%) |
| AdaLN DDPM (best) | 0.243 ± 0.002 (75%) | 0.130 ± 0.002 (99%) |
| AdaLN Flow (best) | 0.234 ± 0.002 (72%) | **0.131 ± 0.002 (100%)** |
| XAttn-Direct | **0.267 ± 0.002 (82%)** | 0.126 ± 0.002 (97%) |
| XAttn-Perceiver-p05 | 0.264 ± 0.002 (81%) | 0.127 ± 0.002 (97%) |
| XAttn-Perceiver-p06 | 0.259 ± 0.002 (80%) | 0.127 ± 0.002 (97%) |

### 4.3. Out-of-distribution generalization (TCGA SKCM)

We evaluate zero-shot transfer to TCGA SKCM, which introduces two shifts relative to 10x TuPro: a patient-cohort and tissue-preparation shift, and a GE-source shift (DeepSpot predictions in place of native Visium). Table 3 reports TCGA SKCM test-split FID together with two delta columns: Δ_baseline_ (each GE model versus its sampler-matched no-GE baseline) and the within-model conditioning ablation Δ_OOD_(GE) = FID(UNI) − FID(UNI+GE). Positive values in either Δ mean GE improves FID under shift.

**Table 3.** Zero-shot OOD evaluation on TCGA SKCM (89,576 test spots, 10,000 subsampled at seed 42). TCGA SKCM FID (H0-mini) at *s*_uni_=4.0, *s*_ge_=3.0; lower is better. Δ_baseline_ = FID(no-GE baseline) *−* FID(model) uses the sampler-matched no-GE baseline (PixCell-B for DDPM, PixCell-Flow-B for Flow). Δ_OOD_(GE) = FID(UNI) *−* FID(UNI+GE). For both Δ columns positive means GE conditioning improves FID (opposite sign convention to Table 1’s Δ). Rows are grouped into pretrained ceiling, no-GE baselines, and GE-conditioned models; within the GE block rows are sorted by Δ_baseline_ descending.

| Model | FID | $\Delta_{\text{baseline}}$ | $\Delta_{\text{OOD}}(\text{GE})$ |
| --- | --- | --- | --- |
| PixCell-Pretrained-XL | **252.8** | – | – |
| PixCell-Flow-B | 730.5 | – | – |
| PixCell-B | 747.1 | – | – |
| PixCell-GE-B | **663.8** | **+83.3** | +6.7 |
| PTPL-XAttn-PMA-B | 690.0 | +57.1 | +309.5 |
| XAttn-Direct | 713.6 | +33.5 | +128.3 |
| PixCell-Flow-GE-B-p05 | 755.1 | –24.6 | –14.6 |
| PTPL-AdaLN-B-p06 | 783.3 | –36.2 | –10.5 |
| PTPL-XAttn-PMA-B-p07 | 805.3 | –58.2 | **+377.5** |

**Table 4.**
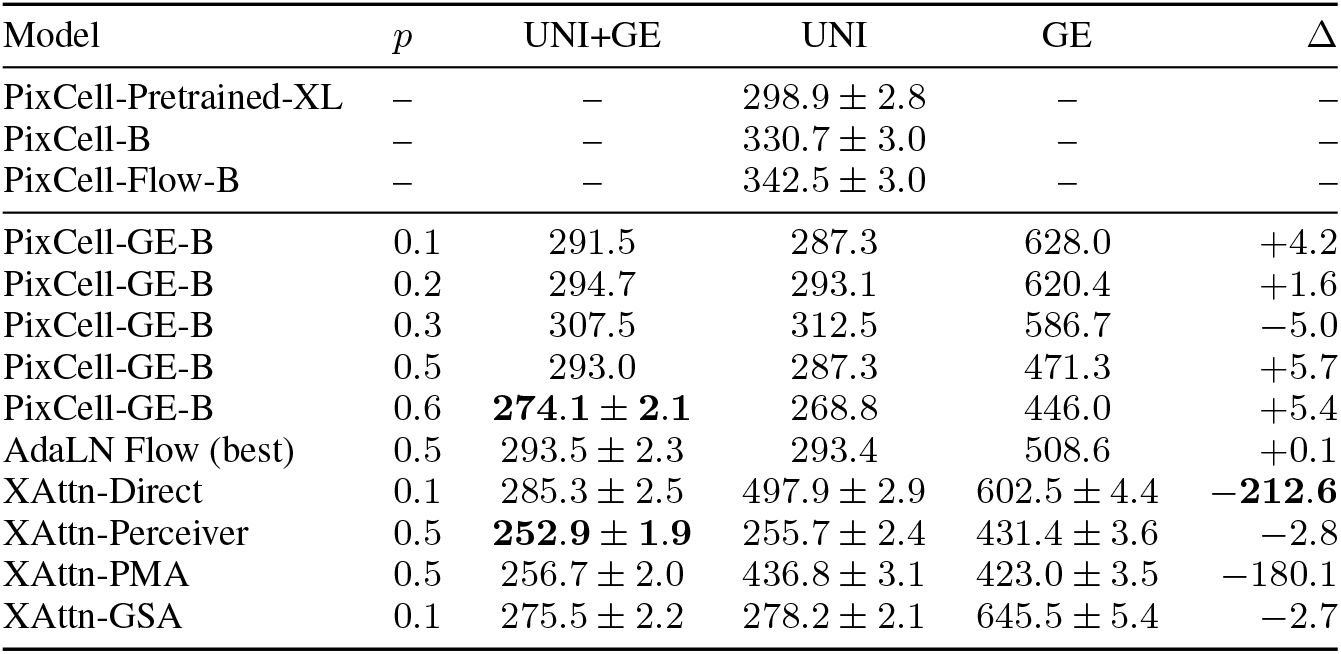
Per-regime test FID. Where a 95% bootstrap CI is available, the cell shows *±*half-width around the point estimate (spotlevel percentile method, *n*=11,776, 1000 resamples). Otherwise the point estimate only. Point estimates match Table 1. Δ = FID(UNI+GE) *−* FID(UNI), where negative values mean GE improves image quality.

| Model | $p$ | UNI+GE | UNI | GE | $\Delta$ |
| --- | --- | --- | --- | --- | --- |
| PixCell-Pretrained-XL | – | – | $298.9 \pm 2.8$ | – | – |
| PixCell-B | – | – | $330.7 \pm 3.0$ | – | – |
| PixCell-Flow-B | – | – | $342.5 \pm 3.0$ | – | – |
| PixCell-GE-B | 0.1 | 291.5 | 287.3 | 628.0 | +4.2 |
| PixCell-GE-B | 0.2 | 294.7 | 293.1 | 620.4 | +1.6 |
| PixCell-GE-B | 0.3 | 307.5 | 312.5 | 586.7 | –5.0 |
| PixCell-GE-B | 0.5 | 293.0 | 287.3 | 471.3 | +5.7 |
| PixCell-GE-B | 0.6 | **274.1 ± 2.1** | 268.8 | 446.0 | +5.4 |
| AdaLN Flow (best) | 0.5 | $293.5 \pm 2.3$ | 293.4 | 508.6 | +0.1 |
| XAttn-Direct | 0.1 | $285.3 \pm 2.5$ | $497.9 \pm 2.9$ | $602.5 \pm 4.4$ | **–212.6** |
| XAttn-Perceiver | 0.5 | **252.9 ± 1.9** | $255.7 \pm 2.4$ | $431.4 \pm 3.6$ | –2.8 |
| XAttn-PMA | 0.5 | $256.7 \pm 2.0$ | $436.8 \pm 3.1$ | $423.0 \pm 3.5$ | –180.1 |
| XAttn-GSA | 0.1 | $275.5 \pm 2.2$ | $278.2 \pm 2.1$ | $645.5 \pm 5.4$ | –2.7 |

Adding GE conditioning to a DDPM backbone improves OOD image quality even when the GE pathway is largely passive at inference: PixCell-GE-B beats the no-GE PixCell-B baseline by 83 FID points (Table 3, Δ_baseline_ = +83.3) while showing only a +6.7 within-model GE-ablation effect. TCGA SKCM carries no native Visium readout, so every test-time GE input is a DeepSpot prediction. PTPL training, which matches this pseudo-label distribution, is the route to H&E-only cohorts and activates the GE pathway substantially under shift. PTPL-XAttn-PMA-B reaches Δ_OOD_ = +309.5 while still beating the baseline by 57 points. Pushing the dropout further amplifies the GE pathway at quality cost: PTPL-XAttn-PMA-B-p07 reaches the panel-leading Δ_OOD_ = +377.5 but trails the baseline by 58 points. Image quality and within-model GE responsiveness on OOD are decoupled. PTPL-AdaLN further transfers across GE sources (Section L).

AdaLN is OOD-neutral across both samplers and both GE sources: the three AdaLN rows in Table 3 (DDPM Visium +6.7, Flow Visium −14.6, DDPM PTPL −10.5) cluster within ±15 of zero. The flat response reflects the fusion mechanism rather than the training signal or the ODE solver.

Data scale dominates every other factor: the PixCell-Pretrained-XL baseline reaches TCGA SKCM FID 252.8, 411 points below every 10x-only model in Table 3 and the only row whose OOD FID is lower than its own 10x TuPro FID, so pan-cancer pretraining and GE-source matching act on orthogonal OOD axes. A tissue-level GE-perturbation analysis on 10x TuPro (Section K, Table 5) reports composites for every GE-conditioned model: composites are pair-selective and bounded in magnitude, and morphological shifts in pixel space remain subtle.

**Table 5.** Tissue-level perturbation composite per source *→* target pair on 10x TuPro (100 reference spots per pair, H0-mini feature space, *λ*_mono_=1.0). Higher composite means the generated tile moves toward the target tissue centroid and does so monotonically in *α*. Three-pair mean averages the three pair-wise medians. ‘–’ = not applicable (no GE input to perturb).

| Model | $p$ | Tumor→Stroma | Tumor→Immune | Immune→Tumor | Three-pair mean |
| --- | --- | --- | --- | --- | --- |
| PixCell-B (control) | – | – | – | – | – |
| PixCell-GE-B | 0.1 | −0.781 | +0.280 | +0.287 | −0.071 |
| PixCell-GE-B-p05 | 0.5 | −0.809 | +0.541 | +0.274 | +0.002 |
| PixCell-GE-B-p06 | 0.6 | −0.159 | +0.547 | +0.620 | **+0.336** |
| PixCell-Flow-GE-B | 0.1 | −0.006 | −0.091 | +0.175 | +0.026 |
| PixCell-Flow-GE-B-p05 | 0.5 | −0.026 | −0.004 | +0.478 | +0.149 |
| PixCell-Flow-GE-B-p05 (UNI-null) | 0.5 | −0.554 | +0.574 | −0.454 | −0.145 |
| XAttn-Direct | 0.1 | −0.108 | +0.282 | −0.678 | −0.168 |
| XAttn-GSA | 0.1 | −0.055 | +0.260 | −0.404 | −0.066 |
| XAttn-Perceiver-p05 | 0.5 | −0.802 | −0.163 | −0.196 | −0.387 |
| XAttn-Perceiver-p06 | 0.6 | −0.050 | +0.448 | +0.057 | +0.152 |
| XAttn-PMA-B | 0.1 | **+0.109** | −0.236 | −0.237 | −0.121 |
| XAttn-PMA-p06 | 0.6 | −0.352 | −0.062 | +0.375 | −0.013 |

## 5. Discussion

### Pseudo-labeled conditioning extends STMDiT to H&E-only cohorts

The PTPL variants realize a predicted-transcriptomics-to-pathology loop: a DeepSpot (Nonchev et al., 2025) predictor supplies denoised gene-expression targets, and the generator produces histology from those predicted profiles. Because DeepSpot pseudo-labels can be computed for any H&E tissue, not only tissues with native Visium annotation, PTPL extends STMDiT to H&E-only cohorts such as TCGA SKCM, where zero-shot evaluation gives a 57-point OOD FID improvement over the no-GE PixCell-B baseline (690.0 vs 747.1) and a within-model GE-conditioning effect of Δ_OOD_(GE) = +309.5 (Section 4.3). This positions pseudo-labeled conditioning as a practical route to virtual tissue synthesis beyond native spatial-transcriptomics coverage, and STMDiT as a tool for hypothesis generation in computational pathology.

### Tissue-level perturbation motivates the next step

The perturbation experiment (Section K) indicates that GE alone does not yet drive strong morphological transitions in our setup. UNI conditioning structurally anchors the morphology when held fixed across *α*, so GE-only intervention produces a directional encoder-space signal but only subtle pixel-space change. Breaking this anchoring through one-sided dropout (GE-anchored training), training-consistent classifier-free guidance, or a different conditioning architecture is a direction we are actively pursuing for the journal version.

Limitations and future directions are discussed in Sections M and N.

## 6. Conclusion

STMDiT demonstrates that jointly conditioning a diffusion transformer on UNI2-h morphology and CancerFoundation gene expression improves both image quality (78-point FID reduction over the no-GE baseline) and transcriptomic fidelity (Oracle AUC reaching 82% of the real-tile ceiling) on 10x TuPro Visium melanoma. PTPL training with DeepSpot pseudo-labels transfers zero-shot to TCGA SKCM (PTPL-XAttn-PMA-B beats the no-GE baseline by 57 FID points; within-model Δ_OOD_ = +309.5), offering a practical route to virtual tissue synthesis beyond native spatial-transcriptomics coverage. CF acts as a frozen projector and the disease-specific gene-to-morphology mapping is learned by the diffusion model on 10x TuPro, so the approach may extend to diseases not in CF’s pretraining.

## A. Data Preprocessing

### A.1. Cohort Details

The Tumor Profiler (TuPro) dataset (Nonchev et al., 2024) comprises 18 H&E-stained tissue slices of metastatic melanoma sequenced on the 10x Visium platform. The slices originate from 9 distinct tissue regions (6.5 × 6.5 mm^2^) across 7 donors, with 2 replicates per region. Each spot was classified into one of five categories (tumor, stroma, normal, lymphoid, blood/necrosis) using histopathology software and subsequently reviewed by a pathologist. Visium captures expression at ∼55 *µ*m spots covering roughly 1 to 10 cells each (Choe et al., 2023).

### A.2. Gene Expression Preprocessing

Raw gene-expression count matrices were concatenated across all 18 tissue slices. Genes expressed in fewer than 10 spots were filtered out. Highly variable genes (HVGs) were identified by selecting the top 3,000 genes using Scanpy (Wolf et al., 2018) with the Seurat v3 flavor, using the sample identifier as the batch key. Counts were then normalized to a target sum of 10,000, log-transformed, and scaled.

#### CancerFoundation embeddings

To obtain gene-expression embeddings, the preprocessed expression profiles were passed through a frozen CancerFoundation encoder (Theus et al., 2024), following the procedures described in the model’s public repository. The resulting latent representations serve as the GE conditioning signal for the diffusion model.

#### PTPL pseudo-labels

Raw Visium counts suffer from technical noise due to limited capture efficiency and sequencing depth, which DeepSpot mitigates by aggregating spatial context across neighboring spots, effectively denoising the expression profiles. For the PTPL variant, gene-expression targets were generated using DeepSpot (Nonchev et al., 2025) with default hyperparameters, trained on the 10x TuPro data to predict all protein-coding genes from H&E image patches. DeepSpot is a deep-set neural network that subdivides each Visium spot into sub-spots to capture local morphology and incorporates neighboring spots for spatial context, using a pretrained pathology foundation model for feature extraction.^2^ The DeepSpot predictions replace the raw Visium counts as the GE input to the CancerFoundation encoder. This provides the pseudo-label source used for zero-shot evaluation on TCGA SKCM in Section 4.3. Pseudo-labeling may attenuate biological signal in addition to technical noise, since binarized dropout patterns in sparse transcriptomic data can themselves carry cell-type information (Qiu, 2020). The PTPL variant thus trades one source

### A.3. Spot Filtering

Before feature extraction, we remove low-quality spots using three image-based filters applied to the spot-centered H&E patch: (i) a *foreground filter* that discards spots where fewer than 55% of pixels are non-white (gray-scale threshold 210); (ii) a *blur filter* that removes spots with Laplacian variance below 20, indicating out-of-focus regions; and (iii) an *HSV tissue filter* that requires at least 30% of pixels to fall within tissue-characteristic saturation and value ranges (*S >* 15, *V <* 240). Spots labeled as “Whitespace” or “Mixed” by the pathologist were additionally excluded.

### A.4. Morphology Preprocessing

#### UNI2-h embeddings

Spot-centered image patches were extracted from each H&E-stained tissue slice, with the patch size defined by the Visium spot diameter. Following the default preprocessing of UNI2-h (Chen et al., 2024c; Mahmood Lab, 2024), patches were resized to 224 × 224 pixels and normalized using ImageNet channel-wise mean and standard deviation. Features were extracted from the embedding layer, yielding 1,536-dimensional morphological embeddings per spot.

#### SD3-VAE latents

For the diffusion model’s latent space, spot-centered patches were extracted at 256 × 256 × 3 resolution and encoded using the frozen SD3 VAE (Esser et al., 2024), producing latent representations *z* ∈ ℝ^32*×*32*×*16^.

### A.5. Dataset Split

To prevent leakage across spatially correlated spots, the 18 tissue slices are split at the donor level before any spot-level operations. Six donors (16 slides) form the training pool and one donor (2 slides) is held out for testing. After the spot filters of Section A.3 are applied, the training pool contains 102,912 spots, which we further subdivide 90*/*10 into 92,621 training and 10,291 validation spots. The held-out test donor contributes 11,776 spots, which form the evaluation set used for all FID, AUC, and perturbation results in the main text. Both the Visium and PTPL variants reuse this split verbatim; only the GE target at each spot differs between the two.

## B. Architecture Overview and Block Diagrams

Figure 1 accompanies Section 2.1 with a system-level view of STMDiT at inference. Three frozen encoders — UNI2-h (Mahmood Lab, 2024) for morphology, CancerFoundation (Theus et al., 2024) for transcriptomics, and the SD3 VAE (Esser et al., 2024) for latent decoding — feed a single trainable PixArt-Σ-B denoiser (Chen et al., 2024b;a) that iteratively denoises a Gaussian latent *z*_*T*_ under both conditioning streams. UNI enters the denoiser via per-block crossattention; GE enters via either AdaLN fusion or per-block cross-attention, detailed in Figure 2. The three forward passes used at inference are combined via dual classifier-free guidance (Equation (1)). On 10x TuPro the GE input is a Visium count vector; under the PTPL variant on TCGA SKCM it is a DeepSpot (Nonchev et al., 2025) prediction.

**Figure 1.**
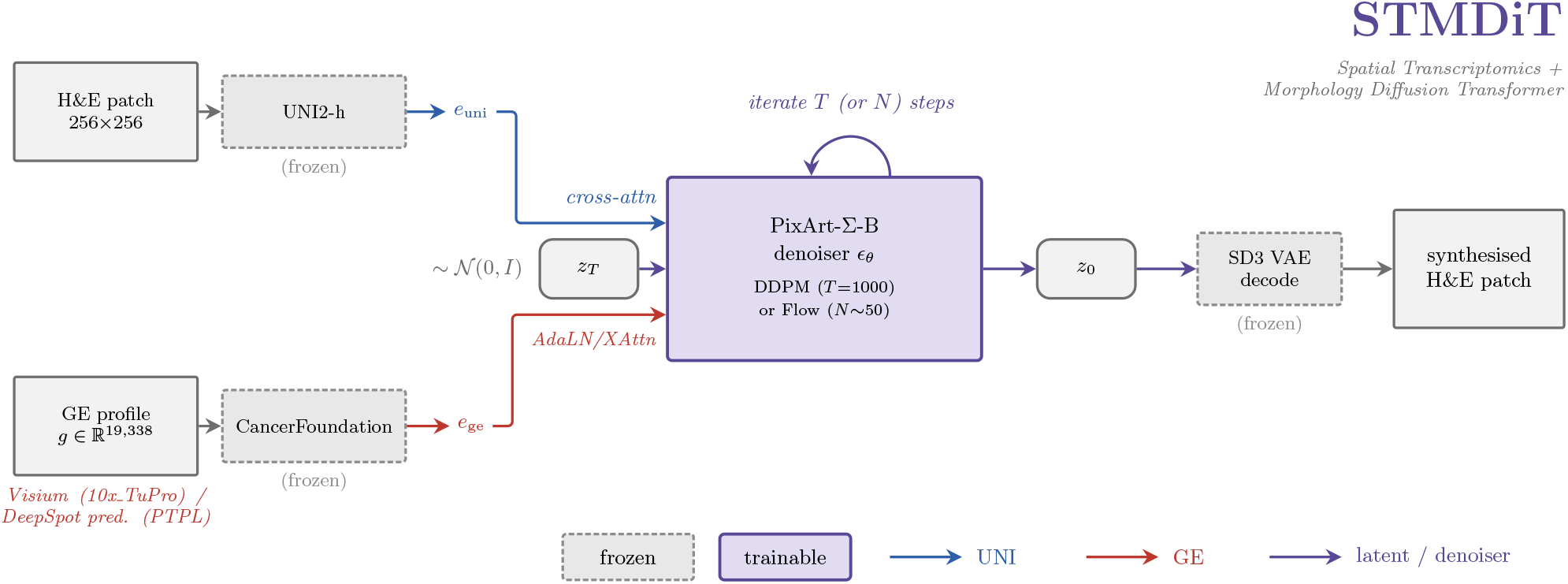
STMDiT inference pipeline. Dashed grey borders mark frozen modules; the solid purple border marks the trainable PixArt-Σ-B denoiser. The two conditioning streams (UNI via cross-attention; GE via AdaLN or extra cross-attention; see Figure 2) and the noise input *z*_*T*_ enter the denoiser at its west edge near the vertical centre. The denoiser is invoked *T* (DDPM) or *N* (Flow) times per sample. The SD3 VAE decodes *z*_0_ into the synthesised 256 *×* 256 H&E patch.

Figure 2 adds a block-level view of the PixArt-B backbone and the two GE-fusion variants we evaluate. Panel (a) shows the unconditioned-on-GE baseline, panel (b) shows AdaLN-GE fusion, and panel (c) shows cross-attention GE fusion.

**Figure 2.**
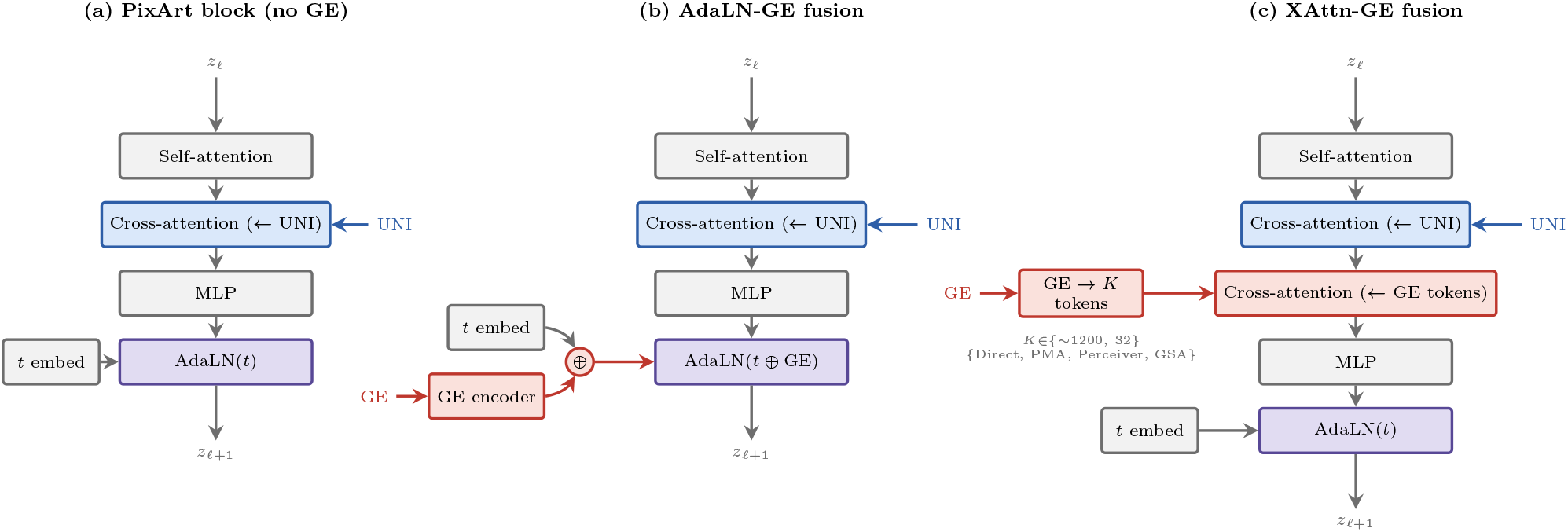
Two ways to add gene expression to the PixArt-B backbone. (a) The base block has UNI cross-attention and AdaLN timestep conditioning. (b) AdaLN-GE fusion combines the GE embedding with the timestep embedding through a learnable gate, so a single AdaLN head modulates every layer on the fused signal. (c) XAttn-GE fusion keeps the timestep AdaLN unchanged and adds a second cross-attention over *K* GE tokens produced by one of four token sources (Direct, PMA, Perceiver, GSA).

## C. Model Family Enumeration

All trained models use the PixArt-B architecture (hidden size 768, depth 12). We train the following families on 10x TuPro.

- **Baselines**: PixCell-B (UNI only, DDPM) and PixCell-Flow-B (UNI only, flow matching (Lipman et al., 2023; Esser et al., 2024)). We additionally include PixCell-Pretrained-XL, a larger pan-cancer PixCell checkpoint, as an external reference.
- **AdaLN**: STMDiT-B with modality dropout *p* ∈ {0.1, 0.2, 0.3, 0.5, 0.6} under the DDPM objective and *p* ∈ {0.1, 0.2, 0.3, 0.5} under the flow objective.
- **Cross-attention**: four token-source variants (Direct, PMA, Perceiver, GSA) at *p*=0.1; Perceiver and PMA are additionally trained at *p* ∈ {0.5, 0.6}.

### Training-data variants: Visium vs PTPL

For three Visium families (AdaLN DDPM, XAttn-PMA, XAttn-Perceiver) we additionally train PTPL counterparts at *p* ∈ {0.5, 0.6, 0.7}. Each PTPL variant conditions on DeepSpot pseudo-labels in place of raw Visium counts, while keeping the same patient-level split, UNI embeddings, and VAE latents. Because DeepSpot pseudo-labels can be computed for any H&E cohort including H&E-only repositories such as TCGA, this enables zero-shot transfer to such cohorts, which we evaluate in Section 4.3. All other hyperparameters are held fixed between matched Visium and PTPL pairs.

## D. Training, Inference, and Evaluation Protocol

This section collects the procedural details required to reproduce every number reported in Section 4. All settings are shared across model families unless explicitly noted.

### D.1. Per-epoch Iteration

Each epoch iterates the full training partition of 92,621 spots once, shuffled with a deterministic seeded generator at the start of the epoch. We do not resample spots within the epoch and do not apply patient-balanced sampling inside the training loop; residual per-slide imbalance is mitigated upstream at data-preparation time by capping the number of retained spots per slide. At effective batch size 128 this corresponds to approximately 723 optimization steps per epoch.

### D.2. Optimization

All PixArt-B models are trained with AdamW (*β*_1_=0.9, *β*_2_=0.999, weight decay 0.01) at a base learning rate of 2 × 10^*−*5^. The schedule is a 1,000-step linear warm-up followed by linear decay to 10% of the base learning rate. Gradients are clipped at a global norm of 0.01, and training runs under <monospace>bf16</monospace>-mixed precision with <monospace>torch.compile</monospace>. We maintain an exponential moving average of the model parameters with decay 0.9999, updated every optimizer step; all evaluation uses the EMA weights. The UNI-only baselines use per-GPU batch size 64 with gradient accumulation of 2 steps; GE-conditioned models use per-GPU batch size 32 with gradient accumulation of 4 steps. Both configurations yield the same effective batch size of 128. Each SLURM submission runs within a 3-day wall-clock budget on a single NVIDIA RTX 4090, and models train for up to 1,000 epochs through chained resubmissions.

### D.3. Inference

Samples for every table and figure in the main text are generated with a DDIM sampler (DDPM-trained models) or the corresponding Euler sampler (flow-matching models), in both cases at 50 denoising steps and a fixed seed of 42. Classifier-free guidance is applied compositionally (Equation (1)), with *s*_uni_ scaling the UNI marginal and *s*_ge_ scaling the GE residual given UNI. We use *s*_uni_=4.0 and *s*_ge_=3.0 at headline inference. Both values were chosen from a joint guidance-scale sweep on FID prior to the main evaluation. The ablation regimes of Table 1 re-use the same scales but zero out the relevant conditioning input during sampling. Inference batches hold 32 samples.

### D.4 Regressor Training

We evaluate transcriptomic fidelity with two independent regressor pipelines to avoid overfitting conclusions to a single encoder family: the regressors differ in backbone, feature reduction, head, and target gene set, so agreement across them indicates regressor-agnostic GE fidelity. Both Oracle and Midnight+MLP regressors are fit on the 92,621 training-split spots and evaluated on the 11,776 test-split spots, reusing the patient-level split of Section A.5; no test-split spot influences regressor fitting. The Oracle regressor reduces the 1,536-dimensional UNI2-h features to 256 principal components and fits one Ridge regressor per gene-count *k* with regularization strength *α* = 100*/*(256 *k*), targeting the top-*k* HVGs by trainingset variance. The Midnight+MLP regressor replaces UNI2-h with the Midnight pathology encoder (Karasikov et al., 2025) (3,072-dim features), feeds those features into a scikit-learn <monospace>MLPRegressor </monospace>with two hidden layers of 1,024 units each, and predicts the full 19,338-gene expression vector after per-spot normalization to 10^4^ counts followed by a log(1 + *x*) transform. Negative predictions are clipped to zero before correlation. Per-gene Pearson *r* is then computed on the held-out test spots, and the integrated AUC in Table 2 averages the resulting curve over *k* ∈ {50, 100, 200, 500, 1000}.

### D.5. Image Quality Metric

Image quality is measured with FID computed on embeddings from the H0-mini pathology encoder (Saillard et al., 2024), a lightweight 768-dimensional distillation of H-optimus-0. For each model we compare FID across the four test-time regimes; a model where GE helps should satisfy FID(UNI+GE) *<* FID(UNI).

### D.6. Bootstrap Protocol

All confidence intervals in Tables 1, 2 and 4 derive from a common spot-level bootstrap with *n*=11,776 and 1,000 resamples. For FID we resample spot indices, recompute H0-mini test statistics on each resample, and report the percentile-method 95% interval; values in Table 4 are written as mean half-width with ± half-width=(upper − lower)*/*2. For AUC we resample spots, recompute pergene Pearson *r*, integrate across the gene-count sweep, and take the percentile-method interval on the resulting AUC distribution.

### D.7. Perturbation Protocol

Tissue-level perturbation samples 100 reference spots per source-target tissue pair and linearly interpolates the input GE embedding between the source and target class centroids at *α* ∈ {0, 0.25, 0.5, 0.75, 1.0}, holding the UNI embedding and the sampler seed fixed across *α*. The composite score sums a transition term (H0-mini distance to the target centroid at *α*=1 relative to the source centroid) and a monotonicity term (negative Spearman correlation of *α* with the per-step distance to the target), with monotonicity weight *λ*_mono_=1.0.

## E. DeepSpot Benchmark on 10x TuPro

To validate that DeepSpot (Nonchev et al., 2025) provides high-quality gene-expression predictions on the 10x TuPro melanoma cohort, we benchmark it against a multi-layer perceptron (MLP) and linear regression. Figure 3 shows mean Pearson correlation as a function of the number of most predictive genes, evaluated via leave-one-patient-out cross-validation. DeepSpot consistently outperforms both baselines across all 19,338 protein-coding genes (AUC = 2732.2 vs. 2158.0 for MLP and 1438.8 for linear regression), confirming that its spatial-context aggregation provides a substantially stronger gene-expression signal than single-spot morphology features alone. This motivates the use of DeepSpot pseudo-labels as the denoised conditioning signal in our PTPL variants.

**Figure 3.**
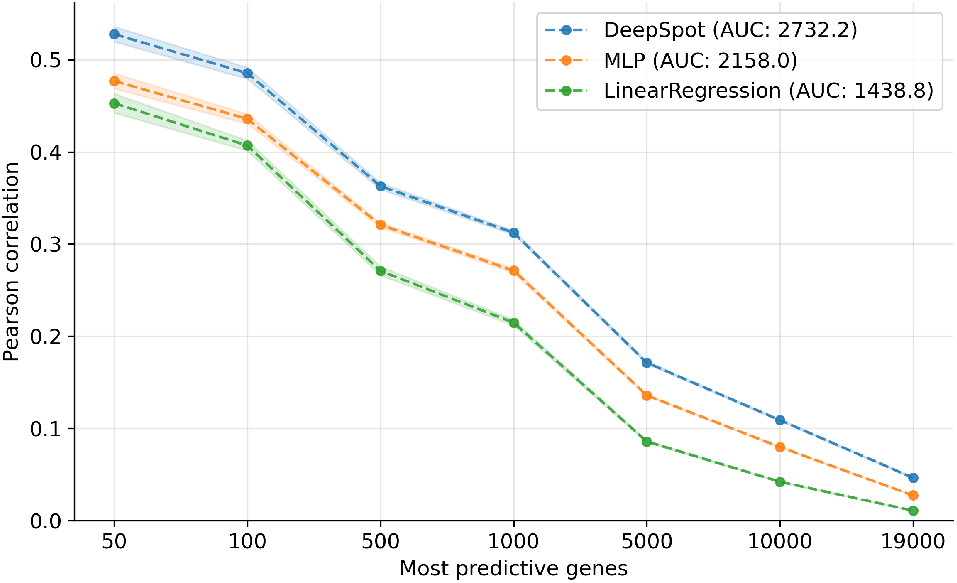
Gene-expression prediction performance on the 10x TuPro dataset under leave-one-patient-out cross-validation. Median Pearson correlation (*±* bootstrap standard error) as a function of the number of most predictive genes. AUC summarizes the area under each curve. DeepSpot outperforms MLP and linear regression across all gene counts.

## F. Additional FID Results

Table 4 extends Table 1 with the full AdaLN DDPM dropout sweep and per-regime FIDs for the cross-attention family.

## G. Cross-Attention Token Source Comparison

## H. Oracle AUC Computation

To establish an upper bound on morphology-based geneexpression predictability, we fit an MLP regressor (two hidden layers of 512 units) on all spots jointly, mapping morphology features to log-normalized expression. This training-set fit (no cross-validation) yields a per-gene oracle Pearson *r*.

## I. Matched-Spot Qualitative Samples

Figure 4 places every ablation model side by side on the same spot key, so a row compares generations of identical morphology across architectures. Rows correspond to the eight GE clusters introduced in Section A and are labelled with their three most distinctive genes. Columns share the reference UNI embedding, the GE vector, the sampler seed, and the guidance scales (*s*_uni_=4.0, *s*_ge_=3.0).

**Figure 4.**
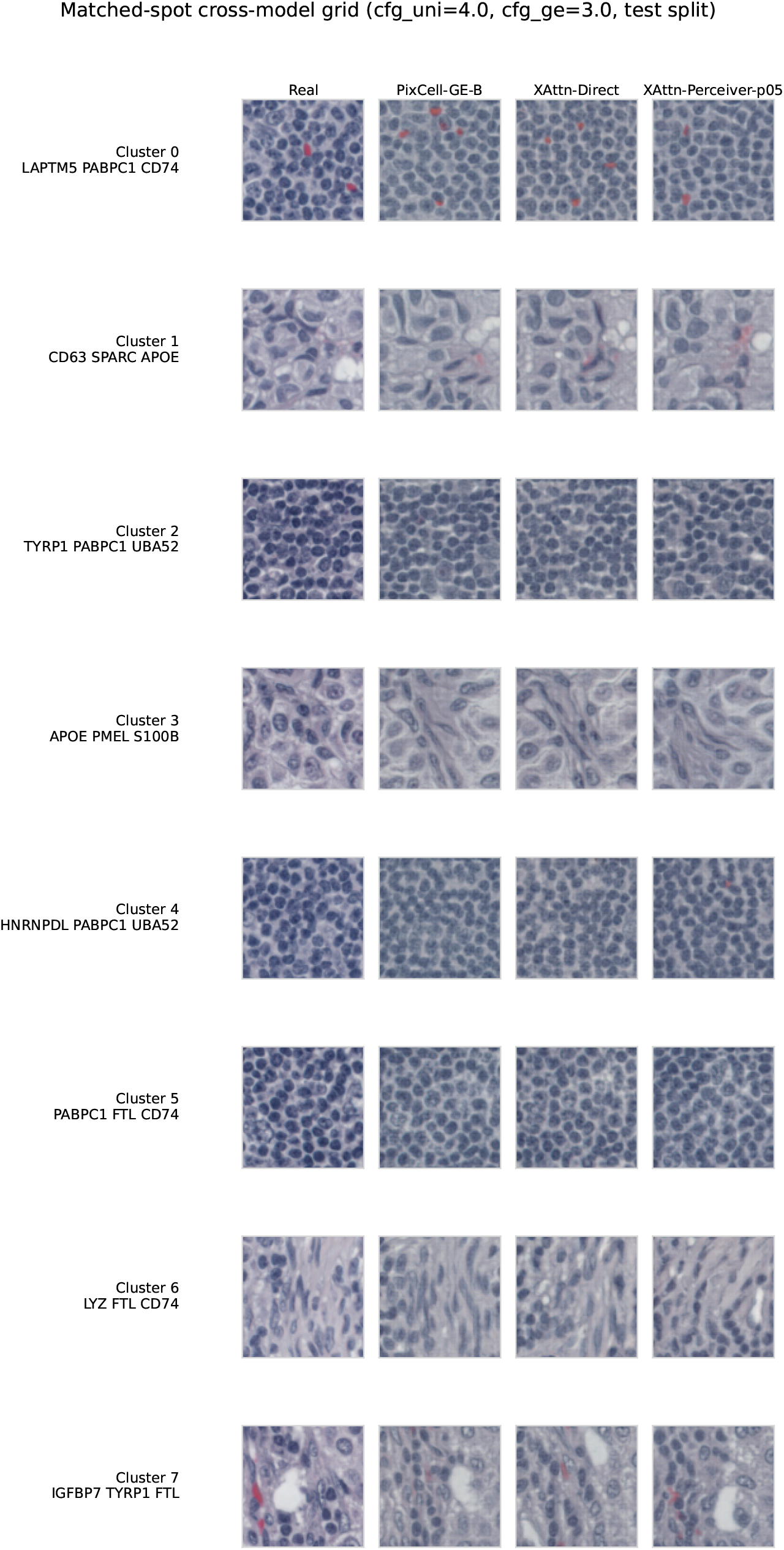
Matched-spot cross-model grid on the 10x TuPro test split. Each row is a single spot rendered by the three GE-conditioned models discussed in Section 4 (PixCell-GE-B AdaLN at *p*=0.6, XAttn-Direct at *p*=0.1, and XAttn-Perceiver at *p*=0.5) alongside the real patch. Rows cover the eight GE clusters and labels list the three most distinctive genes per cluster. All generations use *s*_uni_=4.0, *s*_ge_=3.0, a shared seed, and the same UNI and GE embeddings, so differences across columns reflect architecture alone.

## J. Per-Tissue Quality Visualizations

The spots in 10x TuPro carry pathologist tissue annotations with soft per-class probabilities. We use these labels to audit image quality at a finer granularity than a single dataset-level FID and to compare the three best models at a tissue level. Exemplars are drawn from the test split using a deterministic soft-probability filter that picks high-confidence spots only (argmax agrees with the hard label, *p*_top_ ≥ 0.7, margin ≥ 0.2). The same random seed and the same filter produce the same spot keys across models, so the real reference tile is shared across the model columns in Figure 7.

Figure 5 shows that the best model reproduces the dominant morphology of each tissue class. Tumor tiles carry dense nuclei with irregular stroma, Stroma tiles show fibrous pink matrix with sparse cells, Normal lymphoid tiles show the characteristic small-round-cell population, and Blood and necrosis tiles carry red-cell content and debris. The generated tiles match the dominant staining and texture of their paired real tile without copying it spot-by-spot.

**Figure 5.**
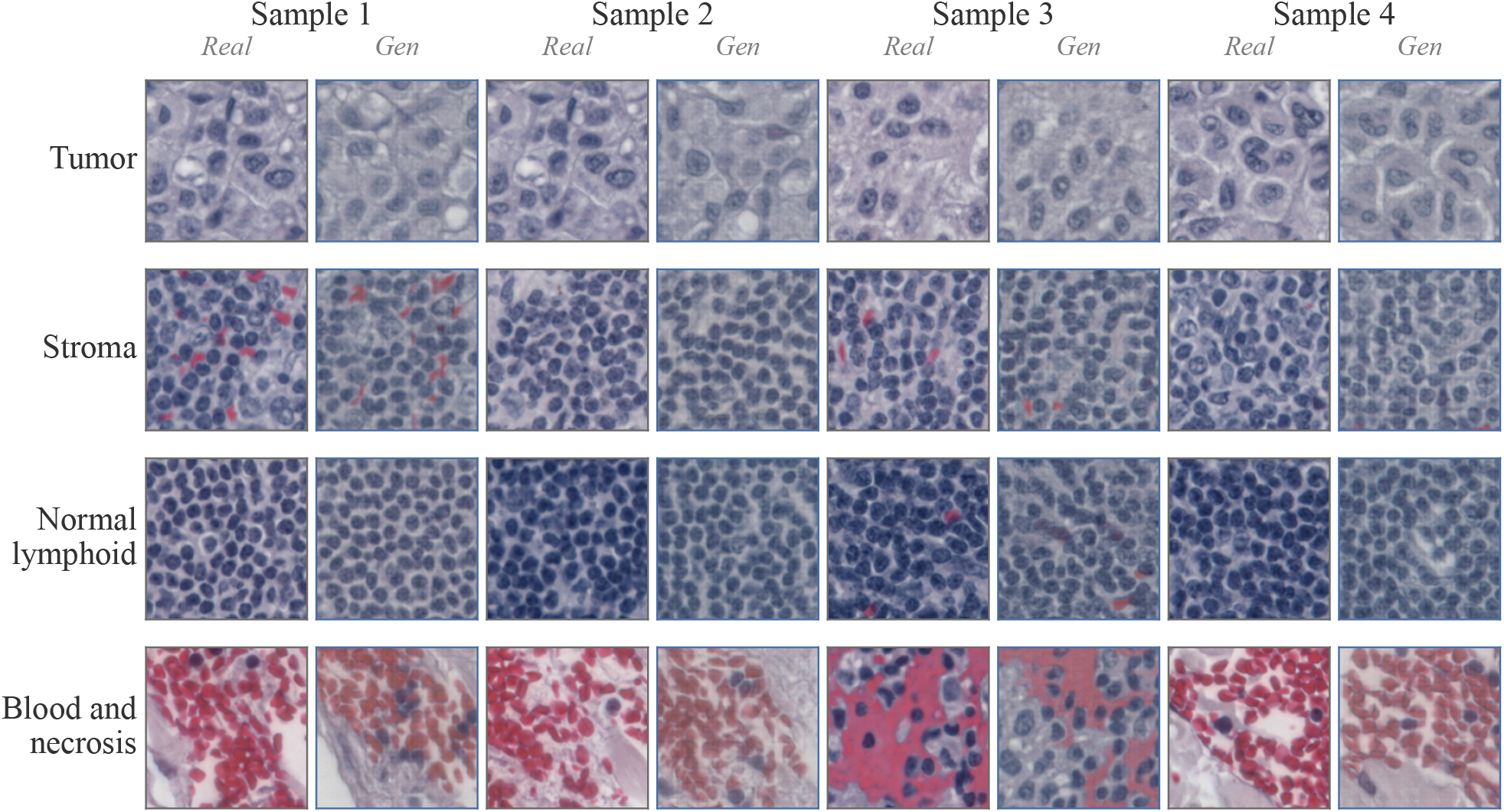
Per-tissue samples from the best model (XAttn-Perceiver at *p*=0.5, FID=252.1). Rows are tissue classes (Tumor, Stroma, Normal lymphoid, Blood and necrosis). Each row shows four *Real* / *Gen* pairs, as labelled above each tile. Spots are sampled from the test split at *s*_uni_=4.0, *s*_ge_=3.0.

Figure 6 reports the per-tissue FID for the same model. Quality is not uniform across classes. The gap between Normal lymphoid (FID=241.7) and Tumor (FID=256.6) is modest, so image quality degrades gracefully on the two common classes. Stroma trails by another 20 points, consistent with the lower spot count (*n*=1741 test-split spots) and the more variable pink fibrous texture. Blood and necrosis (FID=409.2) is the outlier. Its small test-split spot count (*n*=121) inflates the FID estimate and reduces its reliability, so we interpret this bar as an upper bound on how hard the class is rather than a point estimate.

**Figure 6.**
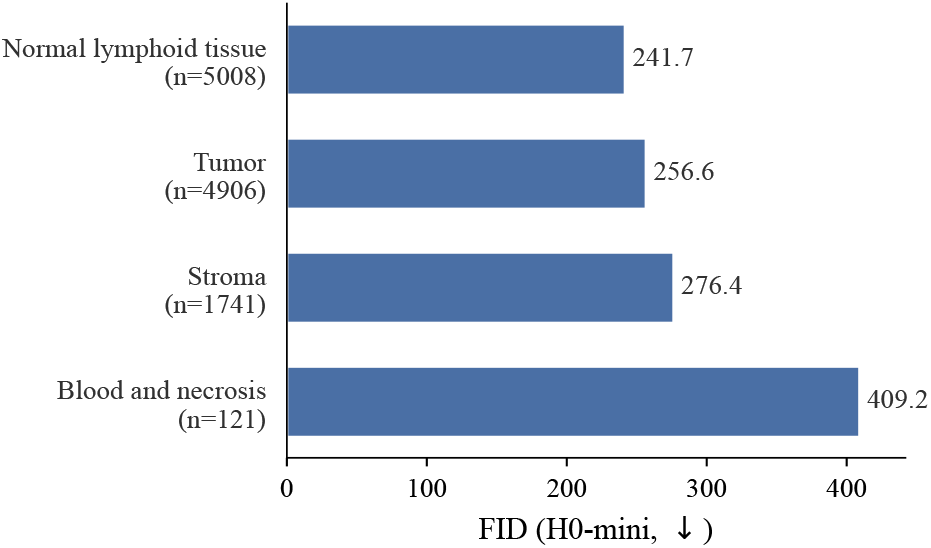
Per-tissue FID for the best model (XAttn-Perceiver at *p*=0.5). FID is computed on H0-mini features over the test-split Visium spots of each tissue class (spot count *n* in the axis label). Normal lymphoid is the easiest class and Blood and necrosis the hardest. The Blood and necrosis estimate is the least reliable owing to its small spot count.

Figure 7 compares the three best models on the same reference spots. The three generators produce plausible tiles in every tissue class, with subtle differences in texture smoothness and nuclear contrast. We do not observe a single model that is visibly best across all four classes, consistent with the close FID and AUC margins between these three configurations in Tables 1 and 2. The per-model rows are reproducible from the manifests of the per-model reports and share spot keys across the Blood and necrosis row, where the small pool means all three models draw from the same high-confidence spots.

**Figure 7.**
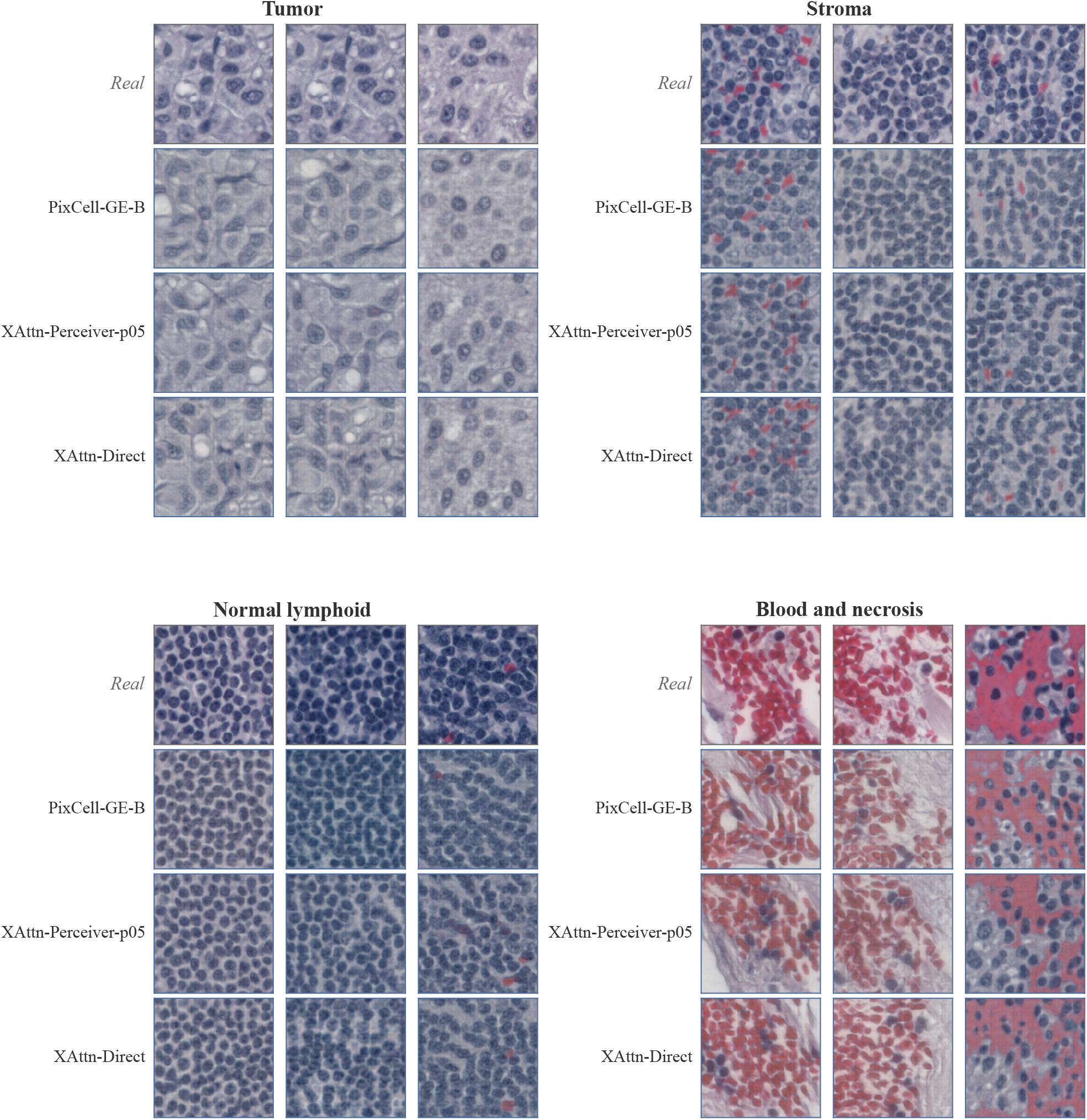
Per-tissue samples across the three best models. Each tissue block shows a shared row of real tiles and one generated row per model (PixCell-GE-B, XAttn-Perceiver at *p*=0.5, XAttn-Direct at *p*=0.1). The real-tile row is shared across models by construction of the soft-probability exemplar filter. All generations use *s*_uni_=4.0, *s*_ge_=3.0.

## K. Tissue-Level Perturbation Panel

We perform tissue-level perturbation by interpolating a model’s input GE profile between the centroids of two tissue classes at *α* ∈ {0, 0.25, 0.5, 0.75, 1.0} and measuring whether the generated morphology follows. Three tissue-class pairs are evaluated (tumor → stroma, tumor → immune, immune → tumor), with 100 reference spots per pair. A composite score combines (i) a *transition* term, the distance in H0-mini feature space that the *α*=1 image moves toward the target centroid relative to the source centroid, and (ii) a *monotonicity* term, the negative Spearman correlation between *α* and distance to target. Higher composite means stronger and more monotonic encoder-space drift toward the target tissue centroid.

Table 5 extends this analysis to every GE-conditioned row whose inference pipeline was run on 10x TuPro. For each source → target tissue pair in Tumor → Stroma, Tumor → Immune, Immune → Tumor we sample 100 reference spots, generate images at *α* ∈ {0, 0.25, 0.5, 0.75, 1.0} with the UNI embedding and the sampler seed held fixed, and report the median composite per pair. The three-pair mean column averages those three medians. The no-GE PixCell-B control and the UNI-null reference for PixCell-Flow-GE-B at *p*=0.5 isolate the contribution of the GE pathway: the no-GE control has no GE input to perturb, so its composite is undefined; the UNI-null row replaces the morphology input with the learned unconditional embedding to test whether GE alone drives the composite.

We report the composite rather than per-*α* tile grids: morphological shifts across *α* are subtle in pixel space, well below the gap between real source and real target tiles, while the composite captures the encoder-space drift quantitatively. High composite values are monotonicity-dominated when transition magnitudes are small: best individual spots reach composite ≈1 with transition near zero, meaning the encoder registers a perfectly ordered but vanishingly small drift toward the target rather than an actual arrival at the target centroid. Combined with the structural anchoring of UNI conditioning across *α*, this is consistent with the bounded composite magnitudes reported below.

Two patterns are visible in Table 5. First, the sign of the composite is pair-specific: several rows that are positive on Tumor → Immune or Immune → Tumor are negative on Tumor → Stroma and vice versa, so a single-pair evaluation can misrepresent a model’s overall controllability. Second, the magnitudes are bounded: no row reaches composite ≥ 0.7 on any individual pair, and the best three-pair mean (+0.336 for PixCell-GE-B at *p*=0.6) is well below the value a perfectly controllable generator would attain. Tumor → Stroma is the hardest transition; XAttn-PMA-B at *p*=0.1 (+0.109) is the only configuration that crosses zero on this pair. Tumor → Immune and Immune → Tumor are more permissive: seven of the twelve GE-conditioned rows are positive on each pair, with peak composites of +0.547 (PixCell-GE-B at *p*=0.6) and +0.620 (same row), respectively. A probable biological reading is that the CancerFoundation encoder, trained on malignant cells, covers lymphoid lineages better than stromal lineages, which would predict easier Tumor ↔ Lymphoid than Tumor → Stroma transitions. Confirming this requires both methodology improvements that rule out alternative causes (model capacity, training procedure, perturbation protocol) and a stromally-enriched encoder, which we leave for follow-up work. No row in Table 5 is positive on every tested pair. We read this as pair-selective directional sensitivity in encoder space: every GE-conditioned family registers a monotonic drift toward the target on a subset of pairs but none does so uniformly across all three.

## L. PTPL distribution sensitivity to GE source

PTPL training matches the conditioning input to DeepSpot’s predicted GE distribution. We probed how much the PTPL OOD result depends on this match by feeding real Visium GE inputs to three PTPL *p*=0.6 variants on the 10x TuPro test split (in-distribution patient cohort; only the GE-source distribution shifts). FID at *n*=11,776 spot bootstrap.

PTPL-AdaLN is essentially robust to GE-source replacement, PTPL-XAttn-Perceiver regresses moderately, and PTPL-XAttn-PMA collapses (Table 6). We did not run the same swap for the *p*=0.5 and *p*=0.7 PMA variants used in Table 3, but the within-architecture pattern at *p*=0.6 suggests their OOD numbers are similarly dependent on GEsource matching.

**Table 6.** PTPL FID under DeepSpot-cond vs Visium-cond inputs on 10x TuPro test split. Lower is better; Δ is the FID regression under input swap.

| Architecture ($p=0.6$) | DeepSpot-cond | Visium-cond | $\Delta$ |
| --- | --- | --- | --- |
| PTPL-AdaLN | 277.5 | 285.6 | +8 |
| PTPL-XAttn-Perceiver | 282.3 | 334.5 | +52 |
| PTPL-XAttn-PMA | 271.2 | **470.9** | **+200** |

## M. Limitations

Our experiments are confined to a single cancer type (metastatic melanoma) and to the PixArt-B backbone. We do not compare directly to the concurrent preprints on gene-conditioned image synthesis (Wu et al., 2024; 2025; Wang et al., 2025; Lohmann et al., 2025), since they target different modalities (gigapixel mouse pup, 3D mouse brain, single-cell Xenium, and mouse kidney Visium HD) and provide neither code nor weights applicable to our Visium melanoma setting. Our Oracle AUC metric also relies on a regressor trained on real spot and gene-expression pairs, and thus inherits any confounding between transcriptomic state and coarse morphological covariates such as tumor content or stromal composition. Dawood et al. (2026) show that H&E-based biomarker predictors can exploit correlated clinicopathological variables as proxies rather than biomarker-specific morphology, so a stratified within-subgroup evaluation is a natural follow-up. The perturbation analysis (Section K) reports observations on a specific training configuration (PixArt-B with a frozen CancerFoundation encoder, single seed per architecture, capped optimization budget). Bounded composite magnitudes and the relative Tumor → Stroma difficulty across architectures could reflect properties of the data, the encoder (CancerFoundation is trained on malignant cells only and may sparsely cover stromal states (Theus et al., 2024)), or artifacts of our specific training procedure such as model capacity, dropout schedule, or random seed. The present analysis does not have the leverage to disentangle these alternatives, so we report the observations without committing to a mechanism. More broadly, the CancerFoundation training corpus is imbalanced across cell lineages (malignant-rich, lymphoid present, stromal sparse), which is an application-dependent limitation of the encoder when the task requires representing or guiding non-malignant cell states.

## N. Future Directions

We plan to scale STMDiT along several axes. Training on larger backbones and multi-cancer ST cohorts such as HEST-1k (Jaume et al., 2024) would test whether dual conditioning generalizes across tissue types and tumor microenvironments. The PTPL pipeline makes it possible to extend STMDiT to large H&E-only repositories such as TCGA, where DeepSpot pseudo-labels can supply the transcriptomic conditioning signal without native ST annotation, opening a path to pan-cancer virtual tissue generation at scale. Emerging single-cell ST platforms (Visium HD, Xenium, CosMx) provide per-cell rather than per-spot expression, which could enable spatially structured gene-expression conditioning instead of the current global-vector approach. Concurrent work on such platforms (Wu et al., 2025; Wang et al., 2025; Lohmann et al., 2025) indicates the potential of this direction. Replacing the CancerFoundation encoder with larger or more general scRNA-seq foundation models, in particular ones with broader coverage of non-malignant tissue and cell lineages, is needed to strengthen the conditioning signal. Our tissue-perturbation results in Table 5, where Tumor → Stroma is the hardest transition across architectures, are consistent with such a gap and motivate pretraining that covers stromal, immune, vascular, and normal epithelial cell populations with density comparable to malignant cells. Beyond encoder coverage, alternative perturbation-modeling formulations and training-time strategies that explicitly account for perturbation responsiveness are an active area of follow-up work.

We also plan to investigate alternative couplings of UNI and GE at inference time. The compositional form of Equation (1) uses three of the four trained regimes, and symmetric or fully factorized variants may redistribute guidance between the two modalities in ways that affect transcriptomic fidelity. More sophisticated dropout schedules (correlated, modality-asymmetric, annealed) could further rebalance the training distribution beyond the single-*p* independent recipe.

## Footnotes

1 GitHub repository: https://github.com/ratschlab/stmdit.

2 Code and pretrained weights: https://github.com/ratschlab/DeepSpot of variability for another.

## Notes

https://github.com/ratschlab/stmdit

### Competing Interest Statement

V.H.K reports being an invited speaker for Sharing Progress in Cancer Care (SPCC) and Indica Labs; advisory board of Takeda; and sponsored research agreements with Roche and IAG, all unrelated to the current study. V.H.K. is a participant in a patent application on the assessment of cancer immunotherapy biomarkers by digital pathology; a patent application on multimodal deep learning for the prediction of recurrence risk in cancer patients, and a patent application on predicting the efficacy of cancer treatment using deep learning all unrelated to the current work. GR is a participant in a patent application on matching cells from different measurement modalities which is not directly related to the current work. Moreover, G.R. is a cofounder of Computomics GmbH, Germany, and one of its shareholders.

